# Parallel evolution at the regulatory base-pair level contributes to mammalian inter-specific differences in polygenic traits

**DOI:** 10.1101/2024.04.19.590285

**Authors:** Alexander S Okamoto, Terence D Capellini

## Abstract

Parallel evolution occurs when distinct lineages with similar ancestral states converge on a new phenotype. Parallel evolution has been well documented at the organ, gene pathway, and amino-acid sequence level but in theory it can also occur at individual nucleotides within non-coding regions. To examine the role of parallel evolution in shaping the biology of mammalian complex traits, we used data on single nucleotide polymorphisms (SNPs) influencing human intraspecific variation to predict trait values in other species for eleven complex traits. We found that the alleles at SNP positions associated with human intraspecific height and red blood cell count variation are associated with interspecific variation in the corresponding traits across mammals. These associations hold for deeper branches of mammalian evolution as well as between strains of collaborative cross mice. While variation in red blood cell count between primates uses both ancient and more recently evolved genomic regions, we found that only primate-specific elements were correlated with primate body size. We show that the SNP positions driving these signals are flanked by conserved sequences, maintain synteny with target genes, and overlap transcription factor binding sites. This work highlights the potential of conserved but tunable regulatory elements to be reused in parallel to facilitate evolutionary adaptation in mammals.

## Main Text

Parallel evolution occurs when two lineages with a similar ancestral phenotype evolve similar adaptive solutions ^1^. Parallel evolution has been shown to occur in natural systems and in experimental evolution experiments at the level of organs, gene regulatory networks, and the amino acid sequences of individual genes ^2–6^. These studies show that the molecular underpinnings of parallel phenotypes are most commonly observed when comparing gene pathways but can also occur at specific amino acid positions ^2^. Parallel evolution in amino acid sequences has been shown for numerous phenotypes in mammals including echolocation ^3^, longevity ^4^, and marine habitat ^3,5,6^. Parallel evolution in non-coding regions has been less well documented, but has been shown to influence coat coloration in beach mice ^7^, pelvic spine evolution in sticklebacks ^8^, and the evolution of flightlessness in birds ^9^. These examples have focused on species with similar selective pressures, but in principle, parallel evolution can occur at orthologous nucleotide positions in non-coding genomic regions of divergent species in response to disparate selective pressures towards similar phenotypic changes ^10^.

While the probability of parallel evolution at any given nucleotide site across the genome is low, complex traits being often highly polygenic – i.e., controlled by thousands of functional genetic variants – therefore present the potential for many sites to be subject to parallel evolution ^11^. This is due in part to the findings that the effect sizes of individual variants underlying complex traits are generally low (i.e., they are not large effect loci) and therefore such functional sites have the possibility to evolve in parallel without drastic effects on phenotypic variation (unlike protein coding variants). For example, human height is controlled by over 12,000 independent genomic regions, with nearly all individually impacting phenotypic variation at best minimally, yet how such variants impact size variation in other species (i.e., in parallel) remains unclear ^12^. In these contexts, long selective sweeps may fix the same variant in divergent lineages so long as that regulatory variant has a similar and advantageous effect on phenotype (see Supplementary Notes 1 & 2).

Though regulatory regions generally evolve more rapidly than protein coding regions and are not limited by the codon-code, promoters can evolve slowly and hundreds of enhancers are conserved in adult mammalian tissues such as the liver ^13^. This suggests that despite substantial regulatory turnover ^14^, thousands of regulatory elements are conserved when all tissues and development timepoints are considered. Within regulatory elements, transcription factor binding preferences are highly conserved in bilaterians ^15^, limiting the potential mutational landscape of regulatory elements at the nucleotide level. This implies that some mutations will have the same phenotypic effect in divergent species. For example, SNP rs12821256 has been shown to alter lymphoid enhancer-biding factor (LEF) transcription factor binding at a KIT ligand (KITLG) gene, contributing to a blonde hair phenotype in humans and a lighter fur coloration when this base pair is experimentally altered in mouse ^16^. Other examples include rs4911178 and rs6060369, which modify hip and knee morphology in humans and predispose individuals to joint disease and similarly alter the morphology of the joint in single base pair replacement mice ^17^. Differential binding of transcription factors (TFs) also can drive gene expression differences between closely related species and deeply divergent lineages. For example, binding of the transcription factor HNF4A is correlated with gene expression in the liver across mouse species ^18^ and some highly-conserved enhancers between zebrafish and mice can drive heterochronic shifts in *Hox* gene expression in transgenic animals leading to dramatic phenotypic outcomes ^19^.

The recent explosion in the number of high-quality human and more generally mammalian genomes and their use in alignment algorithms allows for genome-wide exploration of parallel evolution at the nucleotide level. In this study, we investigated the potential for parallel evolution in the regulation of eleven polygenic traits across mammals using the Zoonoomia alignment of 241 mammalian genomes ^20^ and common single nucleotide polymorphism (SNP) positions associated with those traits in Genome-Wide Association Studies (GWAS) in humans (see Supplementary Note 2). We found that the human alleles at SNP positions associated with height and red blood cell count (RBC) variation are associated with body size and RBC variation between mammalian species. This association holds for deeper branches of mammalian evolution as well as between strains of collaborative cross mice. Furthermore, we provide evidence that the SNP positions driving this signal are flanked by conserved sequences, are in synteny with target genes, and may impact transcription factor binding.

## Results

To evaluate the potential for SNP positions underlying variation in human traits to have evolved in parallel in other mammalian species, we calculated a genotype score for eleven independent complex traits for which we had access to consistent, extensive data across a large number of mammalian clades (See Methods, Table 1). For each trait, we scored >100 species based on their alignment to positions in the human genome significantly associated with variance in that trait by GWAS (see Methods). The ranks of these genotype scores were then tested for a positive relationship with the phenotypic rank of each species for that trait using a phylogenetic least-squares (PGLS) regression. Two traits had a significant positive relationship across mammals: body size (*P* < 1.3e-5; Figure 1A) and Red Blood Cell count (RBC, *P < 3.2e-6;* Figure 2A) (see Table 1, Figures S1-9). Out of the eleven traits investigated, these two traits have the highest heritability (*h*^2^) in the UKBB data. In order to account for bias introduced by the underlying tree structure of the data, the phenotype ranks were randomly assigned to each node using a Brownian model of evolution 1000 times to calculate an empirical *P*-value for each phenotype using the permulation framework ^21^ (see Methods). This resulted in PGLS for both body size (*P* = 0.142) and red blood cell count (*P*=0.074) falling above the standard significance threshold. This suggests that there is a strong phylogenetic component to these associations, i.e., many instances of parallel evolution occurred deeper in the phylogeny rather than between tip species (see below).

**Figure 1.**
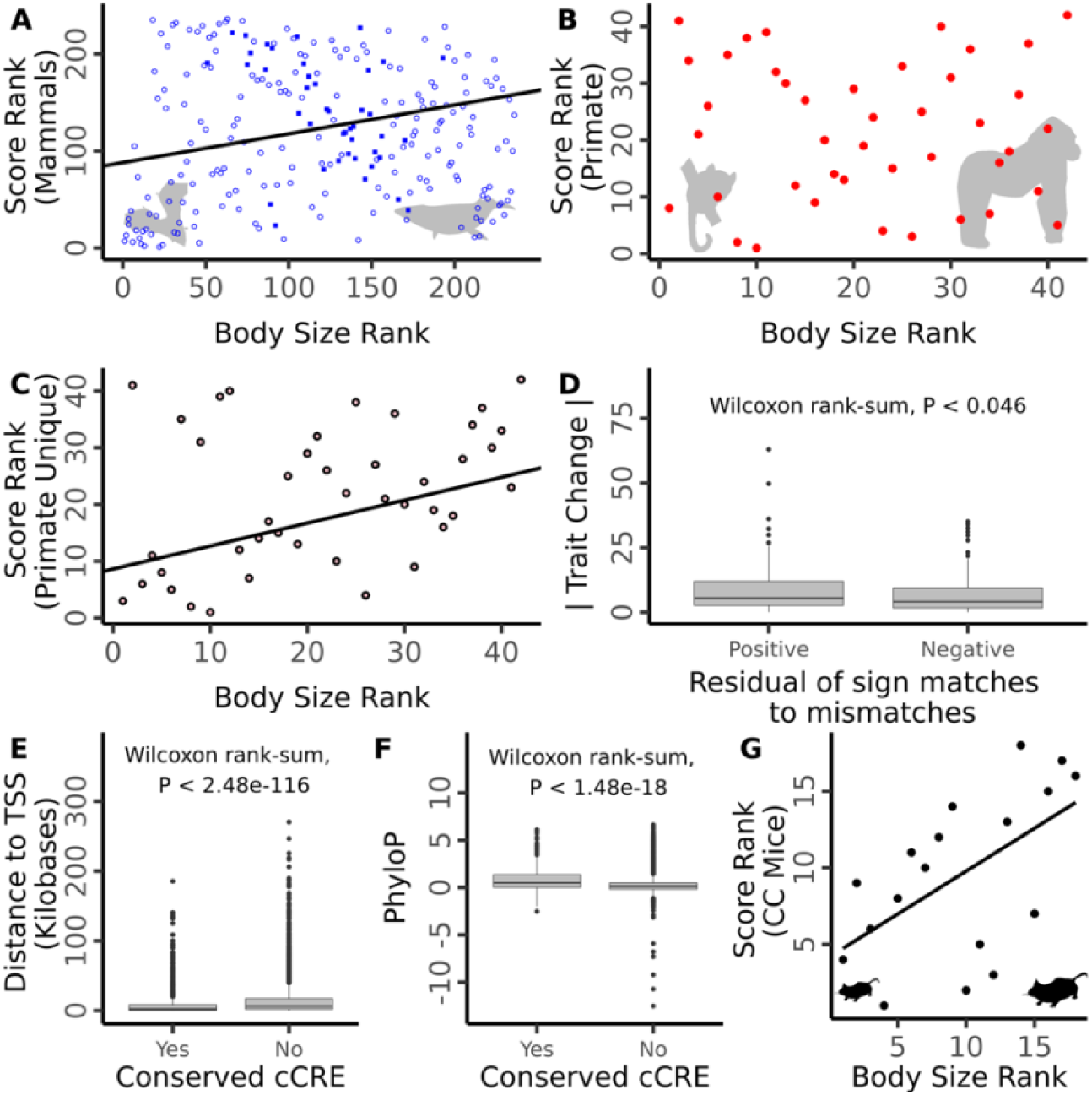
PGLS regression of body size rank against the genomic score computed using mammal conserved SNP positions (A), primate conserved SNP positions (B), or primate unique SNP positions (C). In (A), squares represent primate species, open circle represent non-primate mammals. Silhouettes of the common pipistrelle bat and common minke whale represent small and large mammals, respectively (not to scale). Silhouettes of the gorilla and mouse lemur represent the smallest and larger primates, respectively (not to scale). (D) Comparison of absolute magnitude of body size change for internal branches with a positive residual of matches (allele effect direction matches phenotypic change) to mismatches compared to those with a negative residual. Comparison of the distance to transcriptional start sites (TSS) (E) and PhyloP scores (F) for candidate causative SNP positions which overlap a conserved candidate cis-regulatory versus those which do not. (G) Linear regression of body size rank against genomic score computed for collaborative cross mouse lines. All silhouettes from PhyloPic. Colors of A, B, C as in Figure S10.

**Figure 2.**
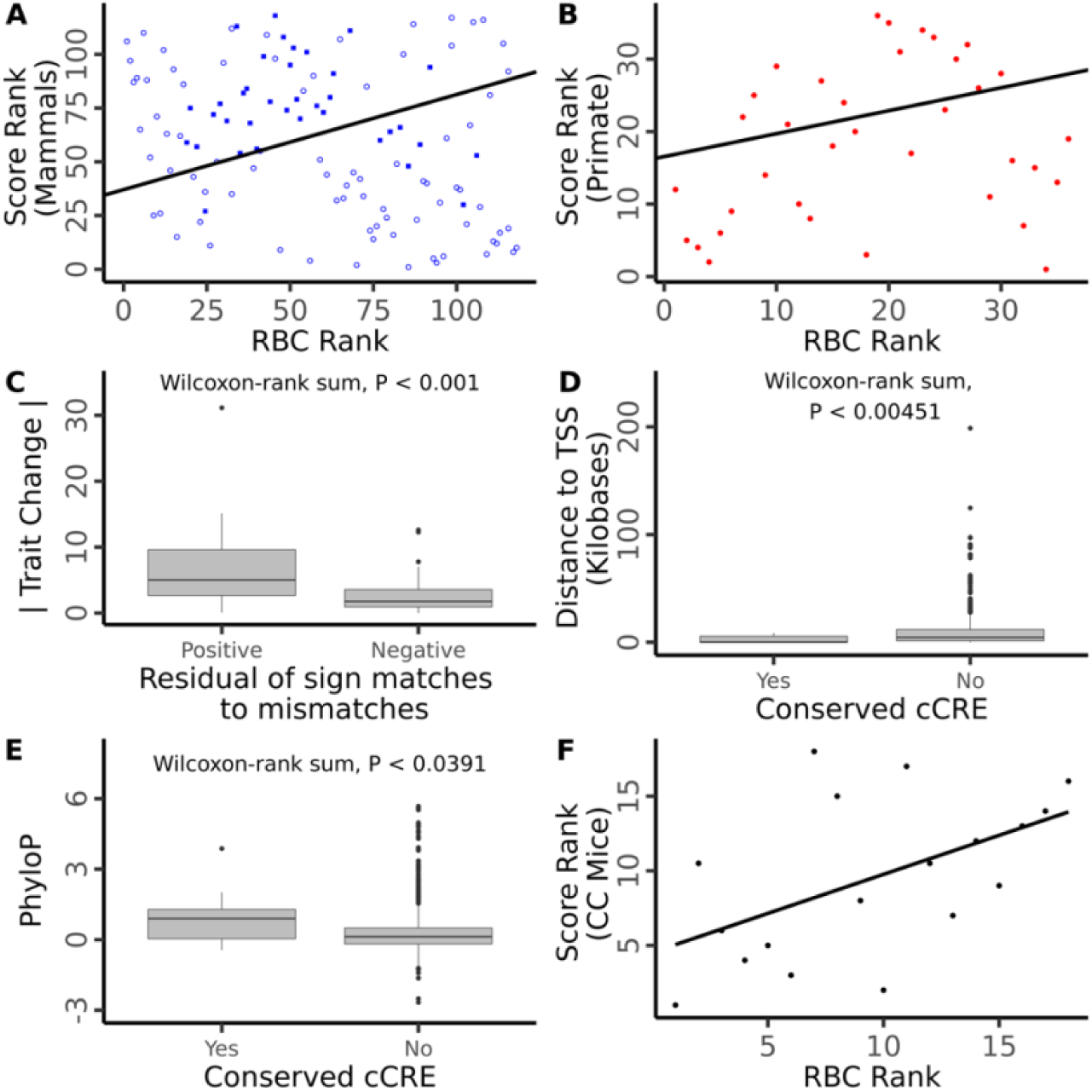
PGLS regression of RBC rank against the genomic score computed using mammal conserved SNP positions (A) or primate conserved SNP positions (B). In A, closed circles represent primate species, open circle represent non-primate mammals. Colors as in Figure 1. (C) Comparison of absolute magnitude of RBC change for internal branches with a positive residual of matches (allele effect direction matches phenotypic change) to mismatches compared to those with a negative residual. Comparison of the distance to transcriptional start sites (TSS) (D) and PhyloP scores (E) for candidate causative SNP positions which overlap a conserved candidate cis-regulatory versus those which do not. (F) Linear regression of RBC rank against genomic score computed for collaborative cross mouse lines.

**Table 1.** Results for the PGLS regressions for eleven traits using genomic scores calculated based on SNP positions conserved in mammals (blue), primates (red), or primates but not other mammals (pink). The number of species (n) for which genomic and phenotypic information is listed for each comparison type along with the number of GWAS significant SNPs, the sign of the relationship, and *P*-values determined by PGLS and after permulations. *P-*values are shown for a Wilcoxon-rank-sum test of enriched trait change along internal branches with positive residuals of trait matches to mismatches and for a Spearman correlation of genomic scores and trait rank in eighteen collaborative cross mice strains. *P < 0.001. Colors as in Figure 1

| Trait | Mammals |  |  |  |  | Primates |  |  |  |  | Primates (Unique) |  |  |  |  | Monophyletic | CC Mice |
| --- | --- | --- | --- | --- | --- | --- | --- | --- | --- | --- | --- | --- | --- | --- | --- | --- | --- |
|  | Species (n) | SNP positions (n) | Sign | PGLS P | Permutations P | Species (n) | SNP positions (n) | Sign | PGLS P | Permutations P | SNP positions (n) | Sign | PGLS P | Permutations P |  | Wilcoxon P | Spearman P |
| Alanine aminotransferase | 115 | 7439 | + | 0.398 | 0.426 | 35 | 12499 | + | 0.147 | 0.135 | 5670 | + | 0.014 | 0.080 |  | 0.112 | 0.229 |
| Aspartate aminotransferase | 115 | 12882 | - | 0.663 | 0.448 | 33 | 22312 | + | 0.395 | 0.297 | 10569 | + | 0.403 | 0.340 |  | 0.189 | 0.319 |
| Calcium | 117 | 6899 | - | * | 0.072 | 35 | 11009 | + | 0.187 | 0.389 | 4647 | + | 0.063 | 0.330 |  | 0.019 | 0.534 |
| Creatinine | 116 | 16365 | - | * | 0.049 | 35 | 26007 | + | 0.452 | 0.357 | 10891 | - | 0.370 | 0.350 |  | 0.484 | 0.703 |
| Eosinophil | 119 | 18877 | + | 0.618 | 0.457 | 34 | 32844 | - | 0.163 | 0.164 | 15556 | - | 0.021 | 0.162 |  | 0.555 | 0.581 |
| Hemoglobin | 120 | 14394 | - | 0.035 | 0.333 | 36 | 25420 | + | 0.574 | 0.369 | 12061 | + | 0.806 | 0.457 |  | 0.859 | 0.531 |
| Lymphocyte | 121 | 12904 | - | 0.036 | 0.192 | 35 | 21808 | - | * | 0.088 | 9938 | - | 0.010 | 0.012 |  | 0.081 | 0.221 |
| Red blood cell | 118 | 18836 | + | * | 0.062 | 36 | 30397 | + | 0.015 | 0.063 | 12948 | + | * | 0.005 |  | 0.001 | 0.013 |
| Body size | 236 | 54451 | + | * | 0.142 | 42 | 87855 | - | 0.472 | 0.290 | 37623 | + | 0.015 | 0.057 |  | 0.046 | 0.008 |
| Urea | 118 | 6119 | + | 0.915 | 0.487 | 35 | 9361 | + | 0.002 | 0.061 | 3684 | + | 0.009 | 0.113 |  | 0.044 | 0.424 |
| White blood cell | 122 | 15176 | - | 0.113 | 0.203 | 36 | 27356 | - | 0.164 | 0.236 | 13449 | - | 0.100 | 0.205 |  | 0.439 | 0.953 |

**Table 2.**
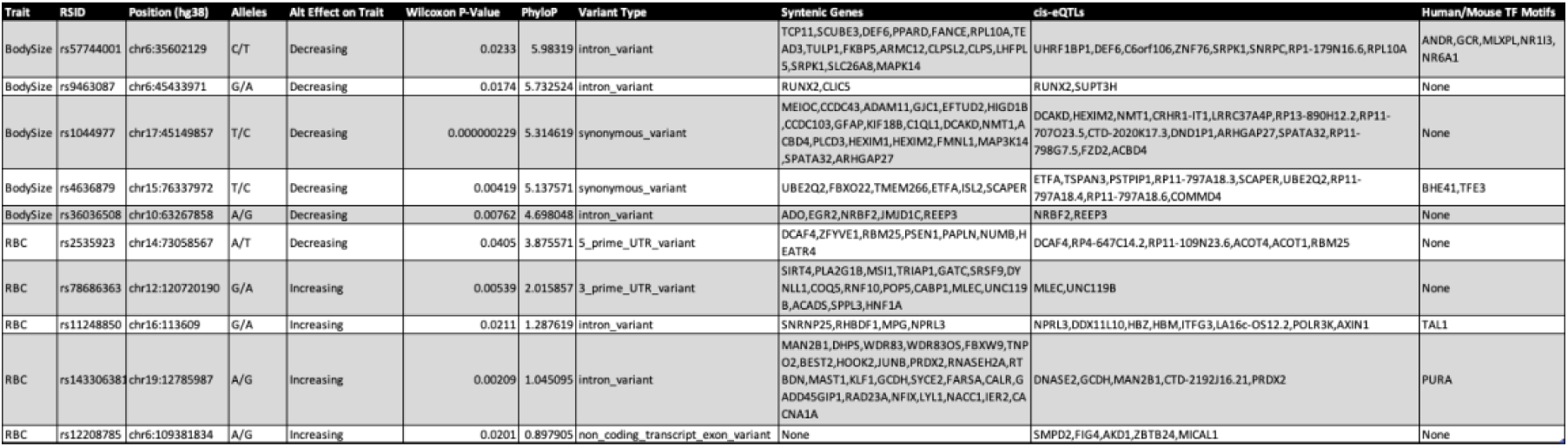
Top candidate causative positions for body size and RBC. Ranked by PhyloP score. Table include columns for the (1) trait, (2) the human RSID, (3) the human genomic position (hg38), (4) the human reference and alternate alleles, (5) the direction of effect of the alternate allele, (6) the Wilcoxon P-value for the association of each allele with the trait across the mammals investigated in this study, (7) the PhyloP score for a range of 21 base pairs centered around the SNP position, (8) the variant type, (9) syntenic genes within a 1 MB window centered on the SNP in human, macaque, dog, cattle, and mouse, (10) genes for which this position is a *cis*-eQTL based on human blood samples, and (11) predicted TF motifs binding around this SNP in both human and mouse.

We next repeated this analysis using just primates since a much larger portion of the human genome aligns to and is conserved only within primates (Figure S10). This approach allowed for more SNPs to be used in computing genotype scores. RBC (*P* < 0.015*, P_permulations_* = 0.063, Figure 2B) and urea concentration (*P* < 0.003*, P_permulations_* = 0.061, Figure S8B) were found to have significant, positive relationships across primates while body size showed no relationship (*P* = 0.47*, P_permulations_* = 0.31; Figure 1B). While mammals overall had a correlation between genotype and phenotype for body size, primates exhibited a strongly negative correlation within this analysis (Figure 1A). When analyzed separately using the larger set of SNPs which aligned within primates, however, primate species showed no relationship between genomic score and body size (Figure 1B), suggesting the larger SNP set has different properties than the smaller set conserved across all mammals. To explore this further, we extracted the SNPs which were alignable in primates but not mammals (henceforth ‘primate unique SNPs’) and reran the PGLS using this reduced set. The primate unique set showed a significant positive relationship for body size (*P* = 0.015*, P_permulations_* = 0.057, Figure 1C), suggesting that sequences specific to the primate lineage might be more relevant to primate body size evolution than more deeply conserved sequences. The primate unique sets PGLS for RBC (*P* = 5.0 e-5*, P_permulations_* = 0.005, Table 1) and alanine aminotransferase (*P* = 0.014*, P_permulations_* = 0.080) also exhibited significant positive relationships.

### Enrichment of Signal in Internal Branches

Since closely related species are often phenotypically similar, many of the genetic changes underlying that shared trait likely became fixed on the branch leading to their last common ancestor. Therefore, if parallel evolution occurs between species as shown in the previous analyses, internal branches with high trait change should be enriched for nucleotide substitutions which drive a corresponding trait change in humans. To address this hypothesis, we identified all the human SNP positions with a fixed substitution in a single mammalian lineage (and not present in any other species) and then tested for an association between an excess of fixed substitutions with the predicted direction of effect and the absolute amount of predicted trait change for each internal tree branch. Using this approach, we found significant associations for both body size (*P* = 0.046, one-sided Wilcoxon rank-sum, Figure 1D) and RBC (*P* < 0.001, one-sided Wilcoxon rank-sum, Figure 2D), as well as for calcium (*P* = 0.019, one-sided Wilcoxon rank-sum, Figure S3D) and urea (*P* = 0.044, one-sided Wilcoxon rank-sum, Figure S8D).

### Identification of Putatively Causative Positions

To identify potential causative variants underlying parallel evolution, we sought to determine which SNP positions underly the correlation between species traits and genome-predicted trait values using PGLS. To do this, we used a Wilcoxon rank-sum test to identify positions where species with a given base pair have different phenotype ranks and then tested whether the sign of these positions matches the effect direction as predicted by GWAS for the corresponding trait in humans (see Methods). This resulted in 12,255 and 992 putatively causative variants for body size and RBC, respectively. For body size, there is a slight enrichment (19 positions) in the number of putative causative variants compared to the number of positions with alleles significantly associated with the trait but in the opposite direction to that seen in humans. RBC shows a slightly reduced set of putatively causative variants compared to mismatches (32 less positions). A high number of mismatches in both cases is expected due to the strong phylogenetic signal present in the dataset so many of positions will be associated with trait values by chance. To restrict our results to the positions most likely to be biologically meaningful, we filtered the putative causative position sets by overlapping them with candidate *cis*-regulatory regions (cCREs) identified in both human and mouse (see Methods). This filtering step produced candidate causative sets of 323 positions for body size (Table S2) and 9 positions for RBC (Table S3). Compared to the removed positions, these refined sets are significantly closer to transcription start sites (TSS) (Body size, *P* < 2.2 X 10^-16^, RBC, P < 0.005, one-sided Wilcoxon-rank sum, Figures 1E and 2D) and have higher local PhyloP scores (Body size *P* < 2.2 X 10^-16^, RBC, *P* < 0.04, one-sided Wilcoxon rank-sum) for both body size and RBC (Figures 1F & 2E). The mean proximity to the nearest TSS for both refined sets is smaller than the average for all cCRE elements (Body size mean = 4461 base pairs versus ENCODE cCRE mean = 6873, RBC mean = 2773, VISION cCRE mean = 3071), suggesting this enrichment is not fully explained by the cCRE filtering step. If causative positions fall within *cis*-regulatory elements, the positions should remain proximate to the target gene in all species. To test this, we identified all genes with a TSS within 500 kb of the candidate positions in five species with well-annotated genomes: human, mouse, dog, domestic cattle, and the crab-eating macaque. We found that many of our candidate positions were syntenic to a least one gene ortholog in each of these species (RBC 4/9 positions; body size 143/310 positions) (Tables 2, S2-S3). These SNPs were generally closely associated with a gene, sitting either in an intron, or directly up or downstream (Figure S11). Of these positions, many have been previously identified as *cis*-eQTLs in human blood ^22^ for at least one of the syntenic genes (RBC 4/4 positions; body size 94/143 positions). While few SNPs have been functionally tested, one of the identified SNPs, rs11248850, sits within a known enhancer element regulating α-globin genes and has been identified as one of two potential causative alleles for a number of hematic traits including RBC count using a massively parallel reporter assay ^23^.

Switching between the allelic variants at each of these positions should result in differential binding of transcription factor (TF) homologs in all species. Since high-quality TF binding motifs are only available for human and mouse, we tested the variants and 10 base pairs of flanking sequence on either side in human and mouse genomic backgrounds for predicted differential TF binding. We found that while some positions were predicted to be bound by a least one TF in both species, our variants were only rarely predicted to substantially disrupt TF binding (Tables S2-3). Subtle alterations to TF binding are consistent with the small effect sizes of these variants in humans and can still be biologically relevant.

### Validation Using Mouse Models

If the polymorphisms underlying trait variation in humans can partially explain differences in average trait values across species, then these same polymorphisms should also help contribute to variation within other species. To test this hypothesis, we utilized data from the mouse Collaborative Cross (CC), a project designed to create a population of mice with levels of genetic variation comparable to natural populations, such as humans ^24^. Phenotypic and genotypic data for 18 CC strains were obtained ^25,26^ and these SNPs were filtered by position against the mammals SNP dataset used in this study (see Methods). The ranked genotype score for each CC line was then calculated as described above for all mammals using this reduced set and correlated with the phenotypic rank of each strain using a one-tailed Spearman’s test. The two genotype-phenotypes pairs which were correlated across mammals –body size (*P* = 0.0086, R^2^ = 0.56, Figure 1G), and RBC (*P* = 0.013, R^2^ = 0.52, Figure 2F) – were the only two traits to show a significant, positive relationship in the CC mice. Using this approach, we identified seven putative causative variants for body size and five for RBC in mice. Two body size (rs3743673 and rs6060539) and one RBC (rs2009094) variant were identified as significantly associated with the trait both across mammals and in CC mice. Of the body size positions identified in collaborative cross mice, two fall within QTL intervals for body weight in the large Gough Island mouse ^27^.

One identified body-size variant (rs6060369, C/T) resides in a highly conserved hind limb enhancer element (termed *R4*) in the *Growth Differentiation Factor Five* (*GDF5*) *cis*-regulatory locus. *R4* enhancer loss-of-function in mice significantly decreases tibial length at one year by markedly decreasing *GDF5* expression *in vivo* as well as *in vitro* in human chondrocytes ^28^. In the same direction of effect we find here, the shorter height variant (T allele) in this enhancer significantly decreases *GDF5* expression *in vivo* in mouse chondrocytes from the hind limb using allele-specific expression studies, and significantly decreases reporter gene activation in mouse and human chondrocytes compared to the tall height variant (C allele) ^28,29^. Moreover, the shorter height variant significantly decreases functional *in vivo* binding of a key hind limb transcription factor PITX1 ^28,29^, highly conserved in functionality across vertebrates and critical for both body size and hind limb length ^8,30^.

## Discussion

Evolutionary adaptation can follow myriad genetic paths, but these possibilities are limited by existing genetic architecture (i.e., regulatory elements and transcriptions factors) and variation therein. While large effect variants may initially exhibit the strongest response to selection – as is often the case in domestication ^31^ – long selective sweeps can act on many loci of smaller effect to reach the phenotypic optimum ^32^. Though similar selective pressures will rarely lead to identical molecular changes ^33^, for highly polygenic traits, whose many causative loci blanket the genome, and sufficiently large numbers of species, the signature of deeply shared genetic architecture can be identified.

In this work, we show that some of the alleles which contribute to human intraspecific differences in RBC and body size are correlated with the corresponding traits across mammals. This agrees with previous research showing that while relatively rare against a backdrop of high regulatory turnover, some regulatory elements maintain deeply conserved function within vertebrates ^34^. It makes sense that while some ultra-conserved elements seem intolerant of any variation ^35^, other conserved elements are potentially tunable and can be used over and over again by evolution to drive similar trait changes. The SNP positions we identify as potentially causative fall into this category, exhibiting high local conservation across the alignment, target genes in synteny, and the potential to alter TF binding relevant to the trait. Our findings also highlight the various paths evolution takes to achieve the same phenotype in different lineages. For example, body size evolution across mammals in general seems to utilize different genomic regions than within the primate clade, potentially due to the substantial transposable element-mediated rewiring of the ancestral primate genome and other clade-specific evolutionary constraints and pressures that have manifested since the deep divergence time between primates and other mammalian orders^36^. In contrast, RBC evolution in primates has used both more recently evolved elements as well as deeply conserved ones. As new mammalian genomes become available, future research will be able to identify additional parallelly evolving regions.

Evolution has produced thousands of extant mammals, each with a unique solution to creating a viable organism using a similar set of genetic tools. Conservation scores have proven to be an effective tool for identifying functional genomic elements ^35,37,38^ but variable parts of the genome can be similarly informative ^39,40^. Our study highlights the use of interspecific patterns of variation to understand the shared biology of complex traits across Mammalia. Since linkage disequilibrium patterns are poorly conserved even between humans and chimpanzees ^41^, an important implication of these results is that interspecific variation in human SNP positions can potentially be used to identify causative variants in humans, such as rs6060369 for body size and rs11248850 for RBC.

## Materials and Methods

### Genotype–Phenotype Pair Identification

GWAS traits used in the UK BioBank were filtered for high heritability (*h^2^*> 0.1), large sample size (n_eff_ > 300,000), inclusion of both sexes, and high confidence (Tables S1 & S4). Traits were manually filtered to avoid highly redundant traits (e.g., lymphocyte percentage versus lymphocyte count), traits calculated based off other traits (e.g., trunk predicted mass), and traits which are challenging or impossible to evaluate comparatively between species (e.g., tense/high strung).

The resulting GWAS trait list was further reduced to those for which phenotypic data were available for at least 100 species. Phenotypic data for body size primarily came from Pantheria ^42^ but was supplemented by additional sources (see Supplementary Materials and Methods). All other phenotypic data came from the Species360 Zoological Information Management System Expected Test Results Database ^43^, which includes expected blood chemistry values based on hundreds of tests of healthy zoo animals. When multiple options were available for a phenotype (such as automated versus manual counting), the method with the largest species sample size was selected. A final round of manual curation involved a literature review to identify traits with a highly variable underlying composition across species, such as total cholesterol ^44^ and total protein ^45,46^, which were excluded from the analysis. This filtering approach resulted in eleven distinct traits with high quality human GWAS data and a large set of mammalian phenotypic data (Table 1). Standing height GWAS in humans was used as a proxy for overall body size when comparing to other species. Adult body mass data was used instead of head to body lengths because the two are highly correlated (Spearman correlation, *R* = 0.987) within the Pantheria database and body weight data is available for more species, increasing the total sample size for this trait.

### Initial SNP-by-Species Matrix Construction

The positions of the 13,791,467 SNP variants used in the UK BioBank were extracted for the 240 mammal species from the Zoonomia whole-genome alignment ^20^ using custom python scripts to produce a SNP-by-species matrix containing the base pair calls. This dataset was then filtered to remove any SNP positions falling within ENCODE blacklist regions since these regions are known to be problematic across various animal genomes ^47^. Only regions on the human autosomes were used due to the more complicated inheritance patterns and sex-specific differences in utilization of the genetic information on the sex chromosomes. Since many mammal genomes are based on a single individual and no data exists on SNP frequencies within these populations, only positions with an unambiguous single base pair call were retained (>99% of all alignments were unambiguous). For the purposes of this study, we assume that all base pair calls are fixed in that species. Only biallelic human SNPs were retained. Since base pairs in other species which do not match either the human reference or alternate allele cannot be interpreted based on human GWAS summary statistics, all base pairs in the alignment which do not match a human allele were removed. Positions with a fixed base pair across all species were also removed. Lastly, because any potentially causative SNPs must be distinguishable from randomly fixed base pairs in a single lineage, we removed any positions with a single substitution which distinguished a single clade (i.e., all species with that allele were more closely related to each other than any other species).

The resulting matrix was filtered to produce two submatrices with all SNPs that have an alignment of 85% across all primate species or 75% across all mammal species. These thresholds were chosen to balance the need to ensure similar alignment percentages across major taxonomic groups while not discarding SNPs that may have failed to align in some species for artifactual reasons. This produced matrices with 2,598,368 SNPs for all mammals (240 species) and 4,337,809 SNPs for all primates (42 species) (Figure S10).

### Genomic Score Calculation and Trait Association Testing

Due to the substantial variation in prediction accuracy of GWAS polygenic risk scores between ^48^ and within human ancestry groups ^49^, the estimated effect size of individual loci (beta scores) are likely meaningless when applied to other species. Therefore, only the direction of the effect for each allele was considered and fixed at 1 or -1 because signage is highly consistent (>90%) across human ancestry groups ^50^. The genomic score for each species was calculated for all SNP positions in the matrix which have been identified at a genome-wide significance threshold of *P* < 5 X 10^-8^ as the sum of the trait increasing allele matches minus the trait decreasing allele matches. The genomic scores were then converted into ranks and regressed against the trait ranks using a Phylogenetic Generalized Least Squares (PGLS) approach ^51^. Rank conversion was used to standardize the scale of the variables, preventing outliers from disproportionately influencing the regression and reducing the impact of small inaccuracies in trait values. The phylogenetic relationships between the species in the alignment were obtained using the TimeTree database ^52^. Because many traits are associated with phylogeny, we tested the associations identified using standard PGLS by utilizing a permulations approach which maintains the tree topology but redistributes the phenotypes using a Brownian motion model of evolution to ensure similar clustering of traits and generate a more accurate null *P*-value distribution ^21^. This approach was repeated for the mammalian conserved SNP positions, the primate conserved SNP positions, and the primate unique SNP positions (those conserved in primates but not mammals).

To calculate genomic scores in the collaborative cross (CC) mice, a new matrix was generated by subsetting the mammalian conserved SNP matrix to those positions which have the same alleles in the corresponding positions of the mouse genome. These positions were identified by lifting over the human SNP positions to mm10 ^53^ and extracting the corresponding mouse SNP data from the Mouse Phenome Database ^54^. Phenotypic data for 18 lines of CC mice was downloaded from the same source and averaged between the sexes ^25,26,54^. The cross strategy used to make the CC lines prevents the lines from having a clear phylogenetic relationship so a standard linear regression was used instead of PGLS to compare the genomic score and trait ranks for the CC lines. Significance was assessed using a one-tailed Spearman’s test.

### Investigation of Evolution in Internal Branches

To test whether clade-specific substitutions are associated with major trait changes on the ancestral branch for that clade, we identified all SNPs which had a base pair fixed in a particular clade and not present in any other species (“monophyletic SNPs”, excluded in the previous analysis). For each genotype-phenotype pair, the degree of phenotypic change was calculated by subtracting the estimated trait value of the direct ancestor from that of the descendent node at each internal node of the tree. The estimated trait values were generated using the phytools fastANC function ^55^ on the ranked phenotype data. For each internal branch, the monophyletic SNPs were intersected with the corresponding GWAS significant SNP sets (*p* < 5 X 10^-8^) and the SNPs with a fixed allele on the branch which matched a human allele associated with the direction of change observed on that branch were counted. Since the total number of monophyletic SNPs was positively associated with longer branches which tended to have large trait change values, we also counted the number of mismatches where the fixed monophyletic SNP at a GWAS-significant position matched the human allele associated with the opposite direction of the trait change. A linear correlation between the number of matching and mismatched positions for each internal node was calculated and the relationship between a positive or negative residual and absolute amount of trait change on each branch was evaluated using a one-sided Wilcoxon rank-sum test.

### Identification of Putatively Causative Positions

Since only body size and RBC showed a consistent signal, only these two traits were investigated further. To identify the SNP positions which are potentially causative for each trait, we used a Wilcoxon rank-sum test to identify positions where species with the human reference allele have different phenotype ranks compared to those with the human alternative allele and then filtered out the positions where the trait-allele association matches the effect direction as predicted by GWAS for the trait in humans.

The potentially causative SNP position sets were filtered for overlap with candidate *cis*-regulatory elements (cCREs) which are present in both human and mouse. For body size, cCRE sets for human and mouse were obtained from ENCODE ^56^ and the intersection performed using *bedtools intersect*. Since cCRE data related to hematopoiesis has been specifically generated in human and mouse ^57^, the VISION cCRE set was used instead for RBC count, following the same overlap approach. For each trait, the PhyloP scores for each SNP position padded by 10 base pairs on either set were calculated using the Zoonomia PhyloP scores ^37^ and an elevation in average PhyloP score for the positions overlapping a cCRE element compared to those without an overlap was calculated using a one-sided Wilcoxon-rank sum test. Similarly, the minimum distance of each position to the nearest transcriptional start site (TSS) as defined in the refTSS database ^58^ was calculated and the positions overlapping a cCRE element compared to those without an overlap were calculated using a one-sided Wilcoxon-rank sum test.

To determine possible target genes regulated by the identified cCREs, each SNP positions was expanded by 500 kb in both directions and this window was overlapped with gene positions from the NCBI refSeq database for hg38 using *bedtools intersect*. If the positions in conserved cCREs are *cis*-regulatory elements for the same gene in all mammals, the positions should be in synteny with their target gene in all mammalian species. The candidate causative positions were therefore lifted over to the mouse (mm10), crab-eating macaque (macFas5), dog (canFam6), and domestic cattle (bosTau9) genomes using the appropriate UCSC liftover files. The refSeq list of gene positions were downloaded for each species and those associated with each position were determined as described in human above. The resulting gene lists in synteny with a given SNP position in each species were intersected in R. Finally, *cis-*expression quantitative trait locus (*cis-* eQTL) data from human blood samples ^22^ was used to further tie candidate SNP positions to target genes.

To determine whether the variants at these positions have the potential to disrupt TF binding across mammals in these positions in a similar manner, we first used the *motifbreakR* (version 2.14.2) *snps.from.rsid* function ^59^ with information from the packages *SNPlocs.Hsapiens.dbSNP155.GRCh38* and *BSgenome.Hsapiens.UCSC.hg38* to get the human sequence surrounding each SNP. Motif enrichments for each sequence variant were identified using the *motifbreakR* function with the following parameters: *filterp = TRUE, threshold = 1e-4, method = “ic”, bkg = c(A=0.25, C=0.25, G=0.25, T=0.25), BPPARAM = BiocParallel::bpparam()).* This step was repeated with the added parameter *show.neutral = True* to identify all TFs binding to the target region, regardless of any difference in binding between alleles. To identify motif enrichments in mouse, the regions were lifted over from hg38 to mm10 using the R *liftOver* package version 1.22.0 ^60^. Mouse reference sequences were extracted using the *BSgenome.Mmusculus.UCSC.mm10* package (version 1.4.3). If the mouse reference base pair at a given position matched either human allele, the other human allele was used as the alternate allele. If the mouse reference allele does not match either human allele, both alleles were listed as alternative alleles. The PWM matrices for either human or mouse came from HOCOMOCOv10 ^61^. The TFs binding to the variants at a given position in both species were intersected to find candidates with potentially conserved function in both species.

## Acknowledgments

We thank Robert Maier for assistance writing the code and critical feedback. We thank David Reich for providing resources and critical advice on this project. We thank Daniel Richard, Steven Munger, Shamil Sunyaev, Michael Hiller, Nick Patterson, and members of the Capellini Lab for helpful discussions regarding this project. We thank Robyn L. Ball for helping us access the collaborative cross mouse data. ASO was supported by a National Science Foundation Graduate Research Fellowship (DGE-1745303).

## Data availability

Many of the phenotypic data used in this study come from the Zoological Information Management System (ZIMS), an international collaboration and repository for the collection, sharing and analysis of knowledge on biological data from wild and zoo animals. The parent data used in this manuscript is available via a ZIMS subscription. All other phenotypic and genotypic data used in this study is from previously published sources and references are provided in the text and supplementary information.

## Code availability

The code used to perform these analyses and to generate these figures is available at: https://github.com/aokamoto-bio/Parallel-Evolution-Mammalia.

