## Supplemental Materials for "Parallel evolution at the regulatory base-pair level contributes to mammalian inter-specific differences in polygenic traits"

**This PDF file includes:**

Supplementary Notes

Sources of variation

The characteristics of common human variation

Linkage disequilibrium and haplotype conservation

Materials and Methods

Additional Body Mass Data for Zoonomia Species

Supplementary References

Figs S1 to S11

Tables S1 to S4 Captions

**Other supporting materials for this manuscript include the following:**

Table 1 to 2, S1 to S4

### Supplementary Notes

1. **Sources of variation.** If a given genomic position is to evolve in parallel in divergent lineages, the same variation must arise before it can be targeted by natural selection. For closely related species, this may be due to standing variation from the ancestral population as is the case in the parallel evolution of coat color in beach mice [1]. In the absence of standing variation, recurrent evolution at a given position is required. While such an event may be rare, this is unlikely to be limiting over evolutionary timescales. A simple calculation multiplying the current human population by the number of *de novo* mutations per person [2] and dividing by the number of positions in the haploid genome suggests that every possible position should be mutated in at least 100 individuals. While mutations are obviously not evenly distributed across the genome, with the real probability of a given mutation governed by the local mutation rate as well as biochemical limitation such as the increased likelihood of transition to transversion events, over many generations and large population sizes, most variants likely appear at least a few times in each lineage. Mutational hotspots are especially likely to generate the variants necessary for parallel evolution at a specific locus [3].

2. **The characteristics of common human variation.** Common genetic variants in modern humans (defined here as variants with a minor allele frequency > 0.01 in the UK Biobank) have a few key properties which are advantageous for this study. First off, millions of human genomes have been sequenced, providing a vast

catalog of common variants and their frequencies in the target population [4]. Second, since common human variants are found in large numbers of people, they can only be mildly deleterious since mutations that are embryonic lethal or otherwise cause severe disease with high penetrance are extremely unlikely to persist in the population at this frequency threshold. Thirdly, since much of this genetic information is paired with detailed phenotypic data, thousands of SNPs have been associated with phenotypic traits through GWAS. Therefore, by evaluating the position of common human SNPs with phenotypic associations in the genomes of other mammals, there is an elevated chance that the SNP positions are more biologically relevant and tolerant of mutation than random genomic positions.

3. **Linkage disequilibrium and haplotype conservation.** While haplotype structure poses a substantial challenge when comparing humans, LD is rapidly disrupted over evolutionary timescales. For example, humans and chimpanzees only have 125 genomic regions in which the same haplotypes are still segregating [5]. Over longer evolutionary timescales, haplotypes are likely to have been lost due to recombination, fixed in the lineages, or acquired additional mutations to form novel haplotypes. Therefore, if some set of variants in the genomes of humans and chimpanzees have the same biological function, those variants might be more readily identified as causative by comparing across species than within species because linked neutral variants in both species are unlikely to be shared due to divergence in haplotype structure. This argument can be extended to more

distantly related species with the caveat that less variants are expected to have the same biological functions due to increasing regulatory divergence between more distantly related species. Conserved regions are under evolutionary constraint so mutations in these regions are more likely to have functional effects. Indeed, SNPs associated with human diseases in GWAS are 1.37-fold enriched in mammalian constrained regions [6]. Thus, the genomes of other species can potentially be used to identify causative variants in humans that lie within important regulatory regions.

### Extended Materials and Methods

#### Additional Body Mass Data for Zoonomia Species

Species without body mass data in the Pantheria Database [7] were supplemented with the additional information listed below. Where only a body mass range was available, the mean was taken within and between sexes.

1. *Allactaga bullata*, data unavailable
2. *Balaenoptera bonaerensis*, male used since only pregnant female data available, 8,350,000 g [8]
3. *Bos indicus*, 280,000 g [9]
4. *Bos mutus,* 546,250 g [10]
5. *Camelus ferus,* 475,000 g [10]
6. *Canis lupus familiaris,* given the variability of dog breed sizes, dogs were excluded from the analysis.
7. *Capra aegagrus*, 53,750 g [10]
8. *Ceratotherium simum cottoni*, 1,900,000 g [11]
9. *Chlorocebus sabaeus,* 3,975 g [12]
10. *Cricetulus griseus,* 40 g [13]
11. *Crocidura indochinensis*, data unavailable
12. *Ellobius lutescens*, 71 g [13]
13. *Equus przewalskii,* 250,000 g [10]
14. *Eubalaena japonica*, 57,595,750 g [14]

15. *Eulemur flavifrons,* 1,850 g [12]

16. *Fukomys damarensis*, 153.25 g [15]

17. *Galeopterus variegatus,* 1,450 g [16]

18. *Ictidomys tridecemlineatus*, 190 g [15]

19. *Murina feae,* 4.8 g [17]

20. *Mus pahari*, 24 g [13]

21. *Myotis davidii,* 5.95 g [17]

22. *Nannospalax galili, ~N. ehrenbergi,* 162.5 g [13]

23. *Neomonachus schauinslandi,* 204,000 g [18]

24. *Neophocaena asiaeorientalis*, 56,000 g [18]

25. *Piliocolobus tephrosceles*, males only, 9,700 g [12]

26. *Rhinolophus sinicus*, 9.9 g [17]

27. *Spermophilus dauricus,* 223.8 g [15]

28. *Spilogale gracilis,* 492.5 g [19]

29. *Tonatia saurophila*, 28.56 g [20]

30. *Tupaia chinensis*, 125 g [21]

31. *Uropsilus gracilis*, data unavailable

32. *Vicugna pacos*, 44,400 g [22]

### Supplementary Figures

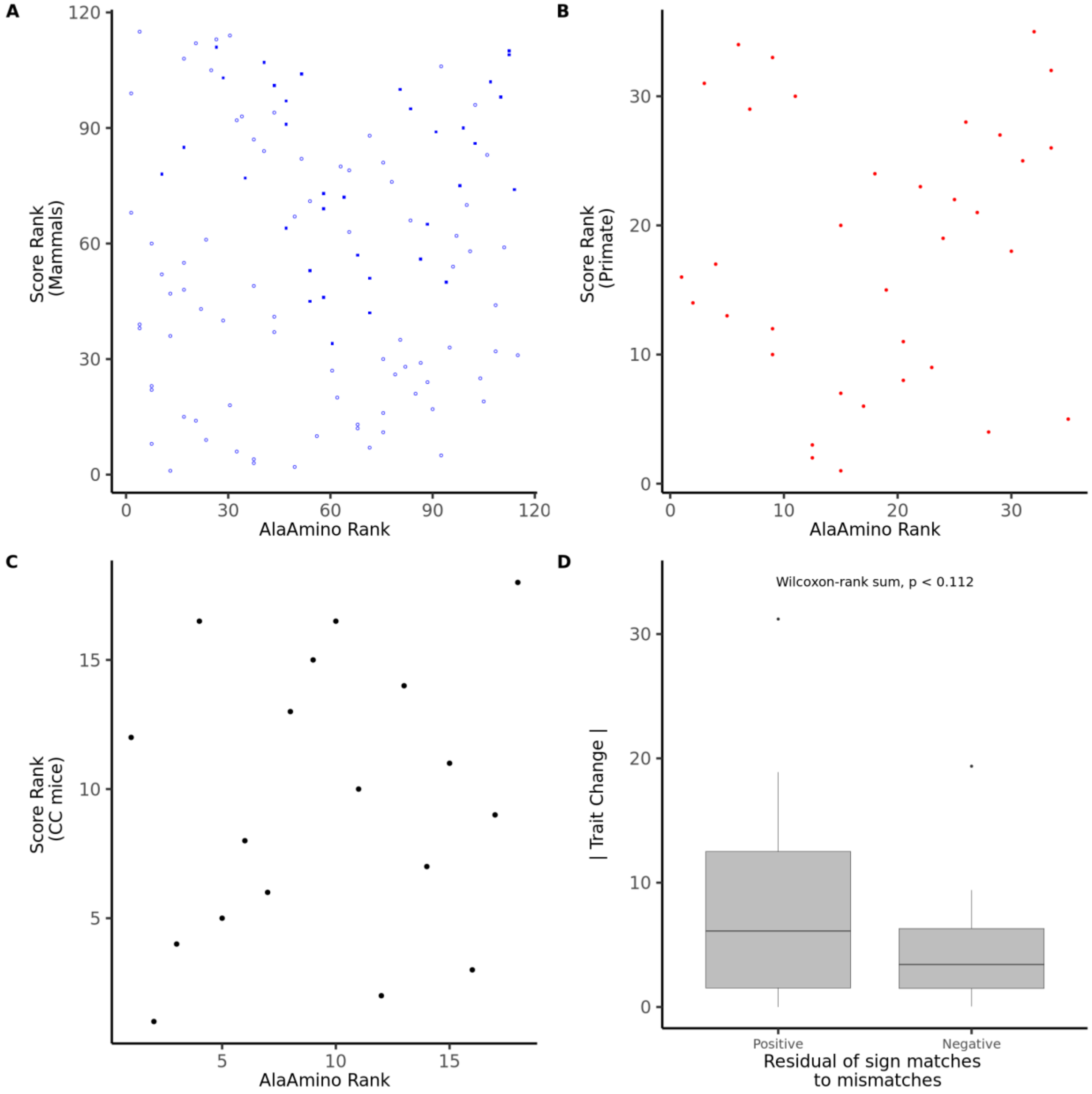


**Figure S1.** PGLS regression of alanine aminotransferase rank against the genomic score computed using mammal conserved SNP positions (A) or primate conserved SNP positions (B). In A, closed squares represent primate species, open circle represent non-primate mammals. (C) Linear regression of alanine aminotransferase rank against genomic score computed for collaborative cross mouse lines. (D) Comparison of absolute magnitude of alanine aminotransferase change for internal branches with a positive residual of matches (allele effect direction matches phenotypic change) to mismatches compared to those with a negative change. Regression lines only show for significant relationships.

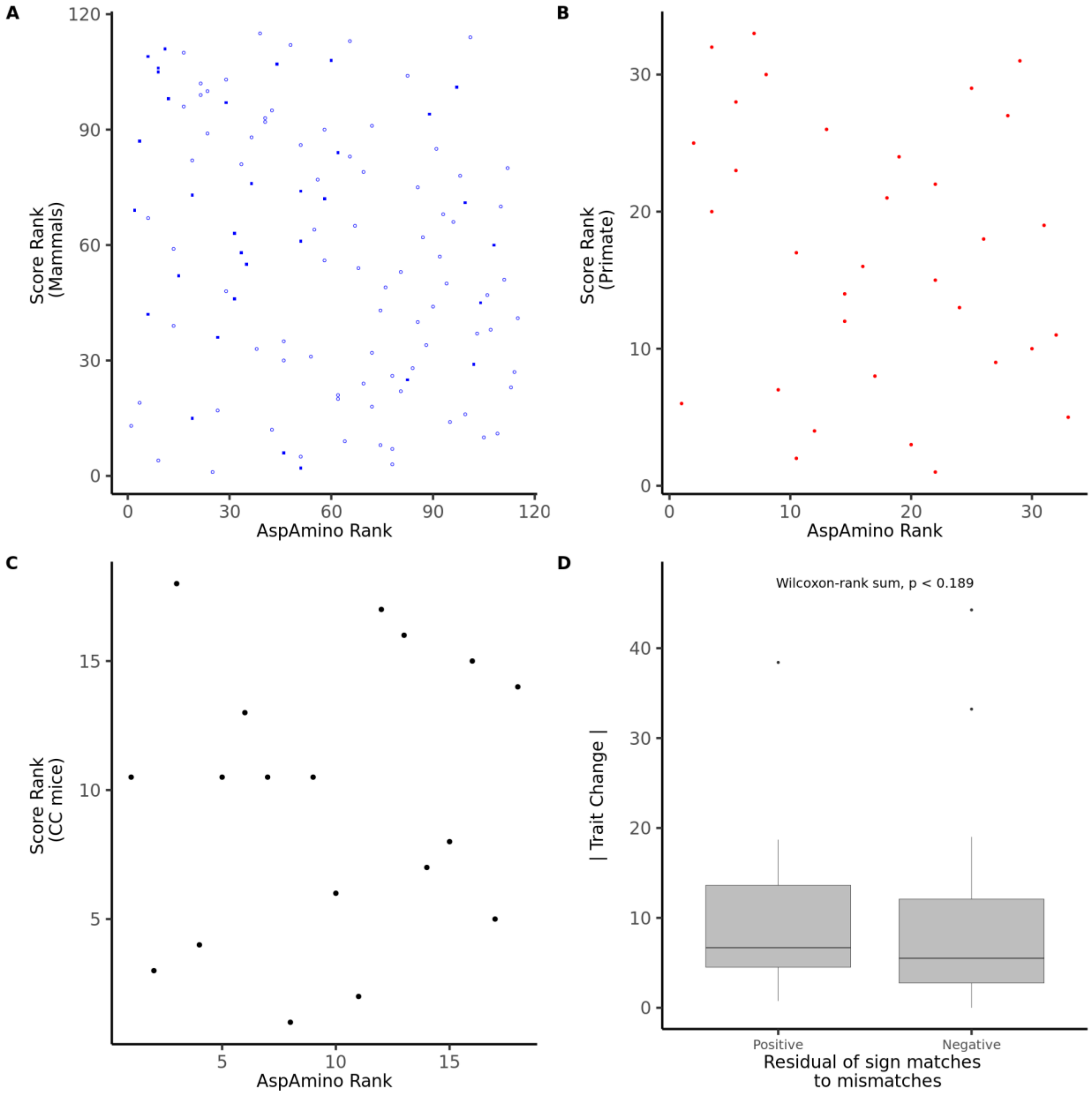


**Figure S2.** PGLS regression of aspartate aminotransferase rank against the genomic score computed using mammal conserved SNP positions (A) or primate conserved SNP positions (B). In A, closed squares represent primate species, open circle represent non-primate mammals. (C) Linear regression of aspartate aminotransferase rank against genomic score computed for collaborative cross mouse lines. (D) Comparison of absolute magnitude of aspartate aminotransferase change for internal branches with a positive residual of matches (allele effect direction matches phenotypic change) to mismatches compared to those with a negative change. Regression lines only show for significant relationships.

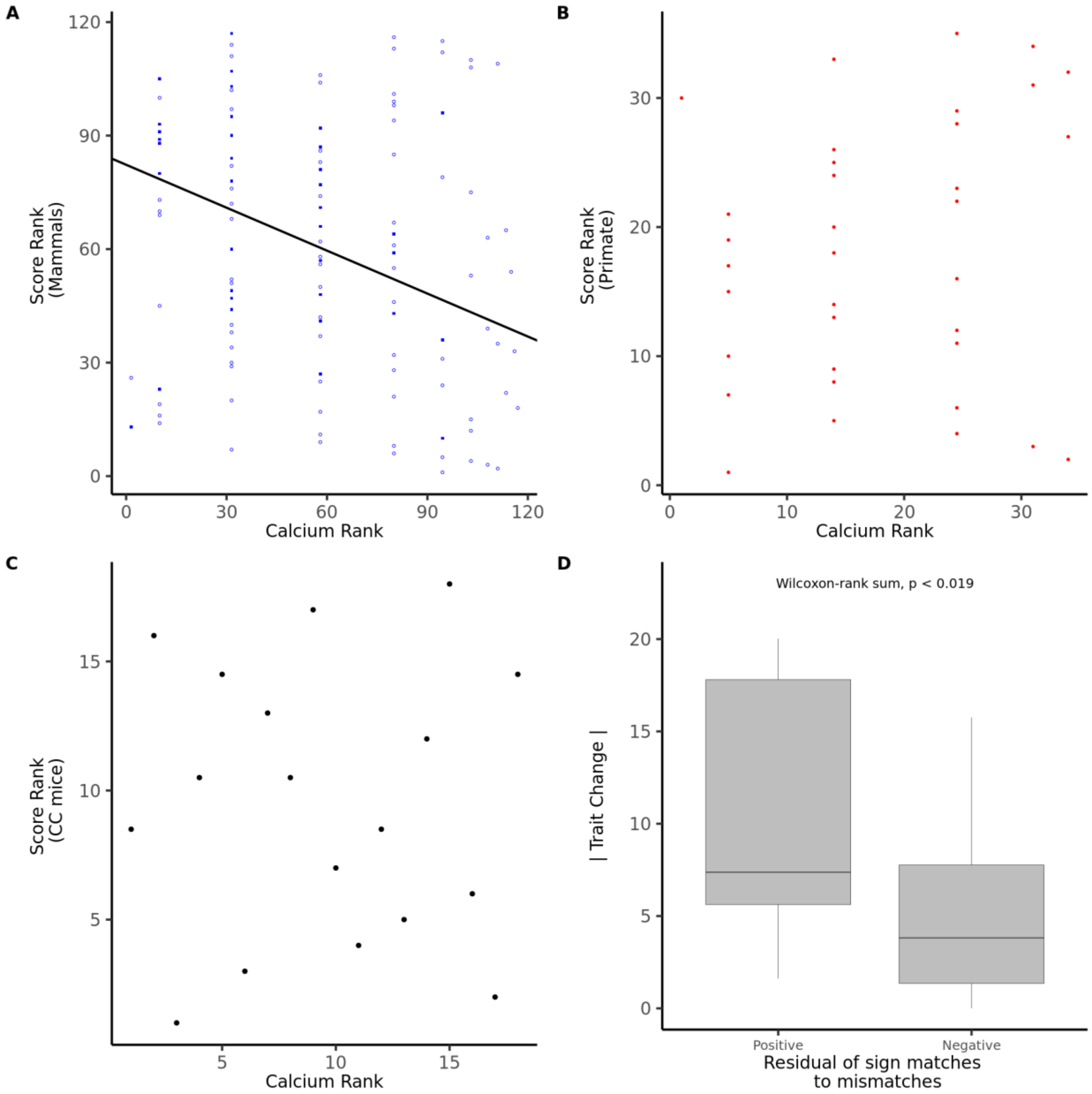


**Figure S3.** PGLS regression of calcium rank against the genomic score computed using mammal conserved SNP positions (A) or primate conserved SNP positions (B). In A, closed squares represent primate species, open circle represent non-primate mammals. (C) Linear regression of calcium rank against genomic score computed for collaborative cross mouse lines. (D) Comparison of absolute magnitude of calcium change for internal branches with a positive residual of matches (allele effect direction matches phenotypic change) to mismatches compared to those with a negative change. Regression lines only show for significant relationships.

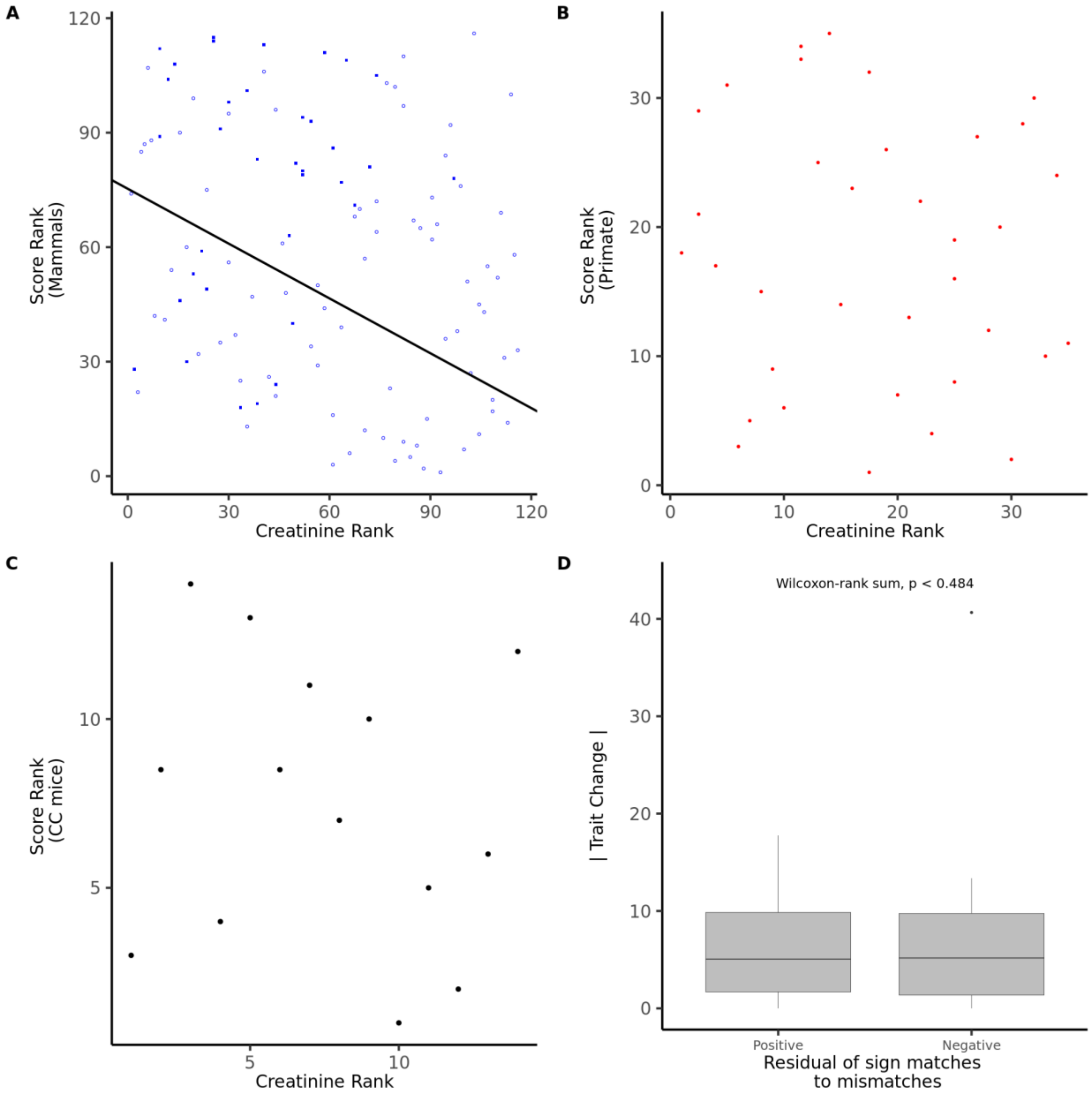


**Figure S4.** PGLS regression of creatinine rank against the genomic score computed using mammal conserved SNP positions (A) or primate conserved SNP positions (B). In A, closed squares represent primate species, open circle represent non-primate mammals. (C) Linear regression of creatinine rank against genomic score computed for collaborative cross mouse lines. (D) Comparison of absolute magnitude of creatinine change for internal branches with a positive residual of matches (allele effect direction matches phenotypic change) to mismatches compared to those with a negative change. Regression lines only show for significant relationships.

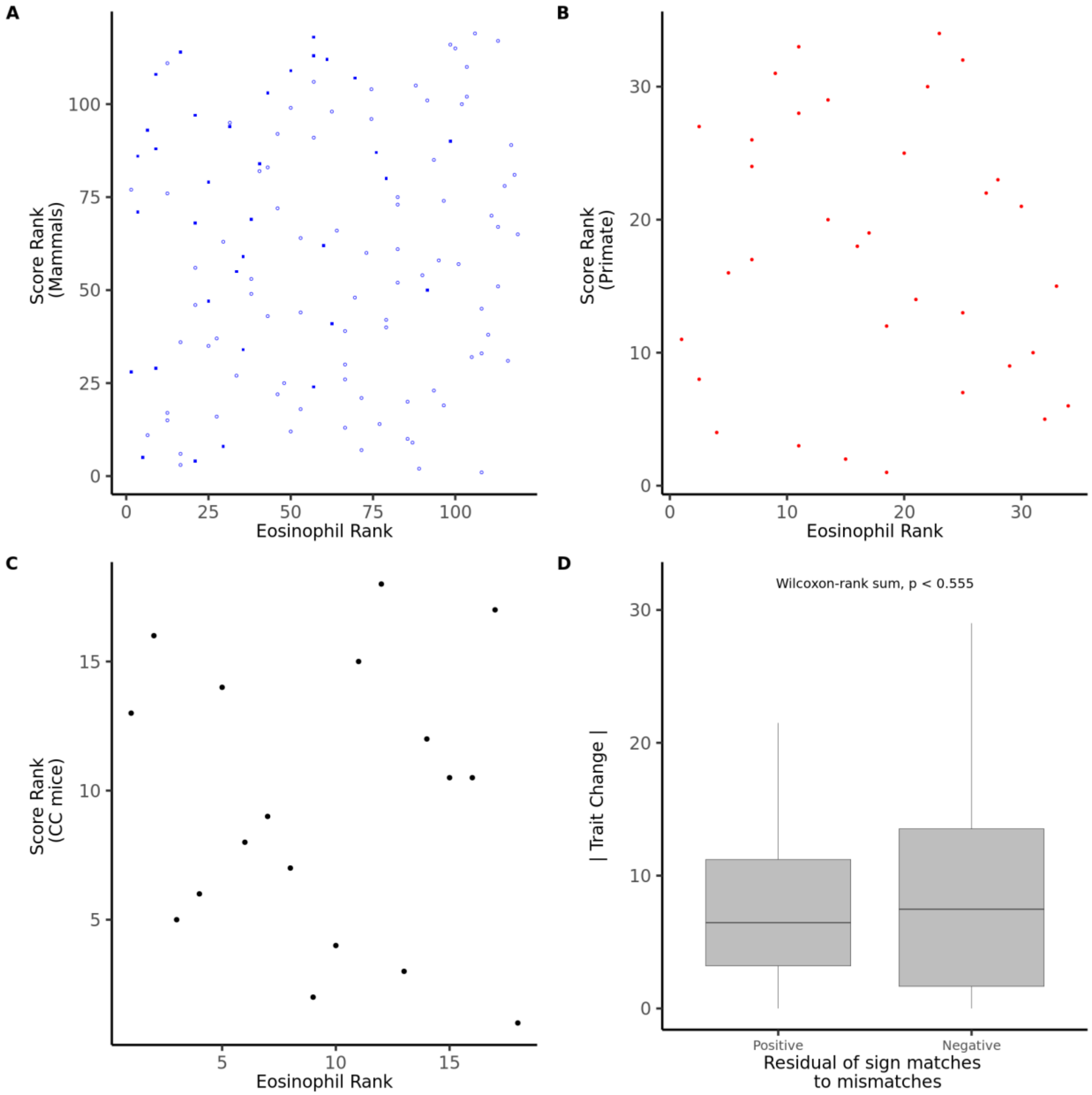


**Figure S5.** PGLS regression of eosinophil rank against the genomic score computed using mammal conserved SNP positions (A) or primate conserved SNP positions (B). In A, closed squares represent primate species, open circle represent non-primate mammals. (C) Linear regression of eosinophil rank against genomic score computed for collaborative cross mouse lines. (D) Comparison of absolute magnitude of eosinophil change for internal branches with a positive residual of matches (allele effect direction matches phenotypic change) to mismatches compared to those with a negative change. Regression lines only show for significant relationships.

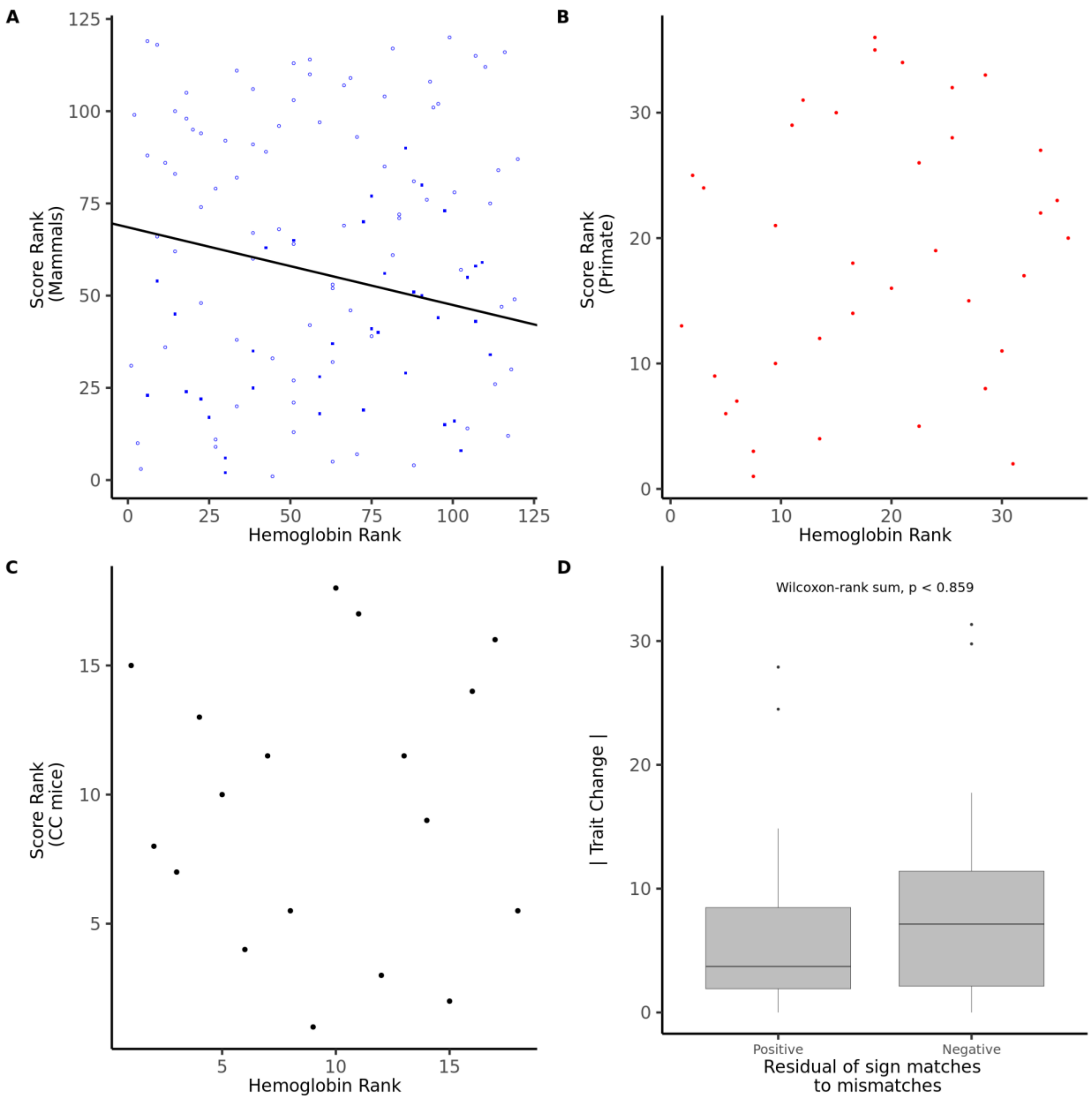


**Figure S6.** PGLS regression of hemoglobin rank against the genomic score computed using mammal conserved SNP positions (A) or primate conserved SNP positions (B). In A, closed squares represent primate species, open circle represent non-primate mammals. (C) Linear regression of hemoglobin rank against genomic score computed for collaborative cross mouse lines. (D) Comparison of absolute magnitude of hemoglobin change for internal branches with a positive residual of matches (allele effect direction matches phenotypic change) to mismatches compared to those with a negative change. Regression lines only show for significant relationships.

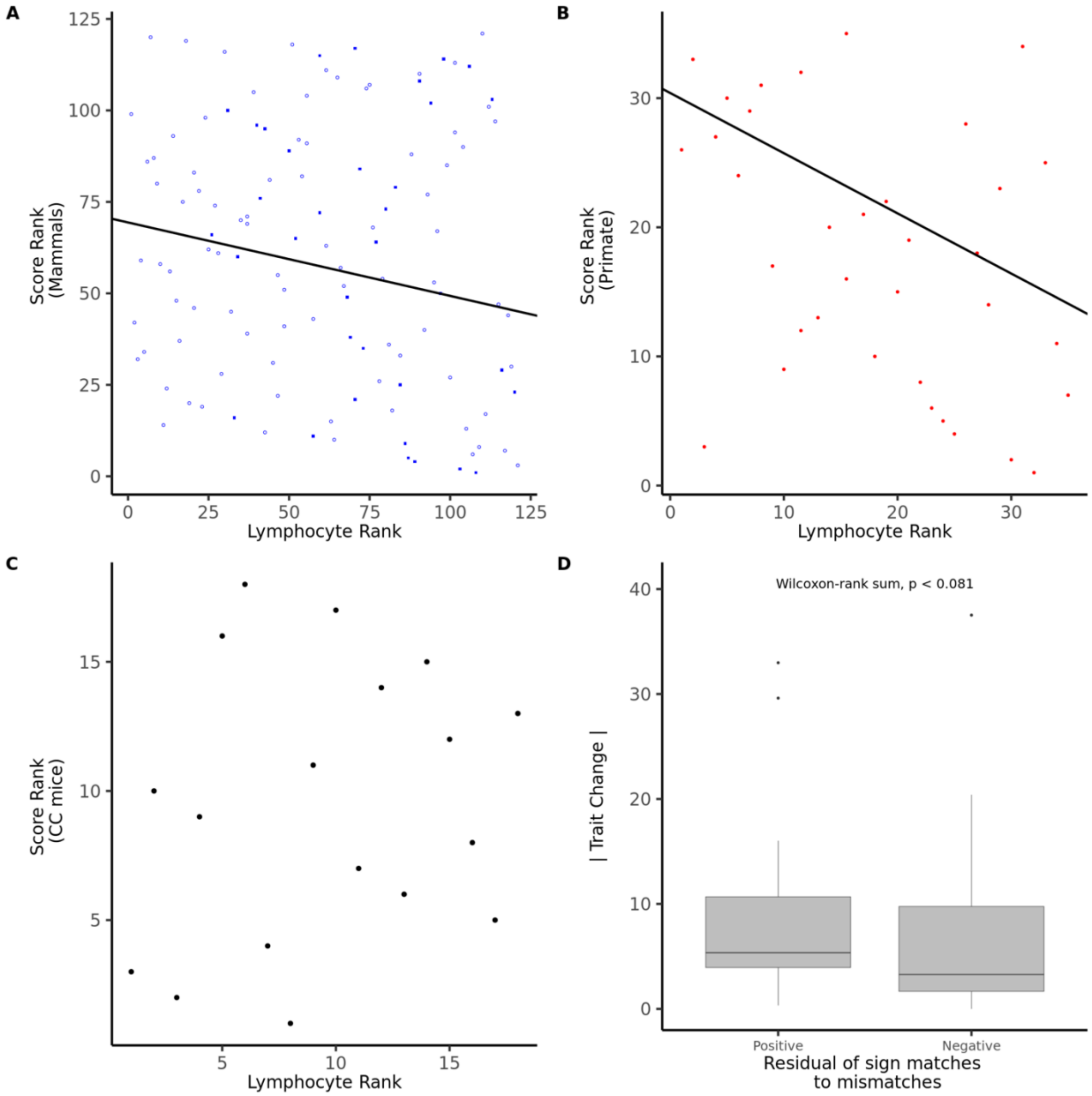


**Figure S7.** PGLS regression of lymphocyte rank against the genomic score computed using mammal conserved SNP positions (A) or primate conserved SNP positions (B). In A, closed squares represent primate species, open circle represent non-primate mammals. (C) Linear regression of lymphocyte rank against genomic score computed for collaborative cross mouse lines. (D) Comparison of absolute magnitude of lymphocyte change for internal branches with a positive residual of matches (allele effect direction matches phenotypic change) to mismatches compared to those with a negative change. Regression lines only show for significant relationships.

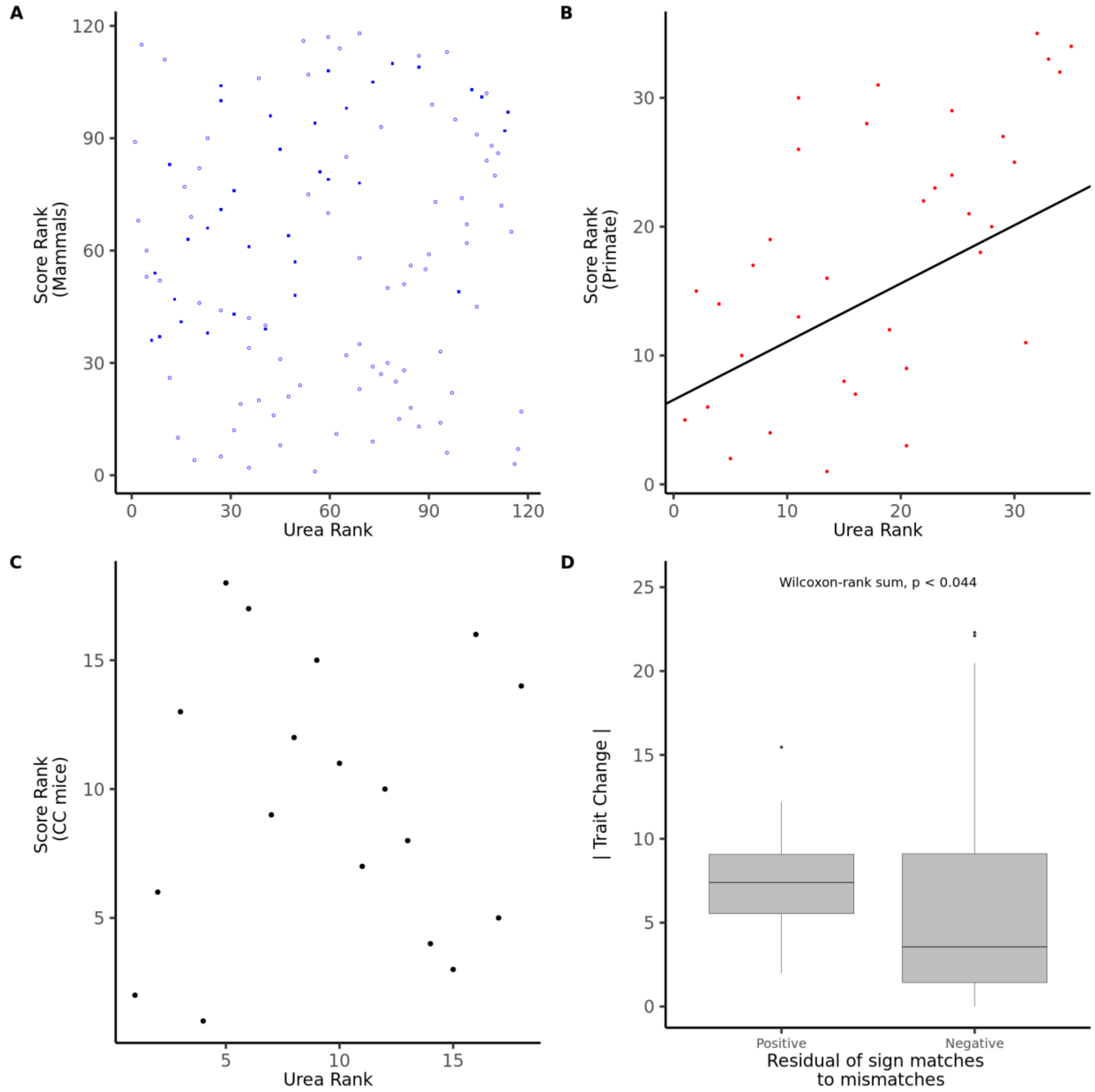


**Figure S8.** PGLS regression of urea rank against the genomic score computed using mammal conserved SNP positions (A) or primate conserved SNP positions (B). In A, closed squares represent primate species, open circle represent non-primate mammals. (C) Linear regression of urea rank against genomic score computed for collaborative cross mouse lines. (D) Comparison of absolute magnitude of urea change for internal branches with a positive residual of matches (allele effect direction matches phenotypic change) to mismatches compared to those with a negative change. Regression lines only show for significant relationships.

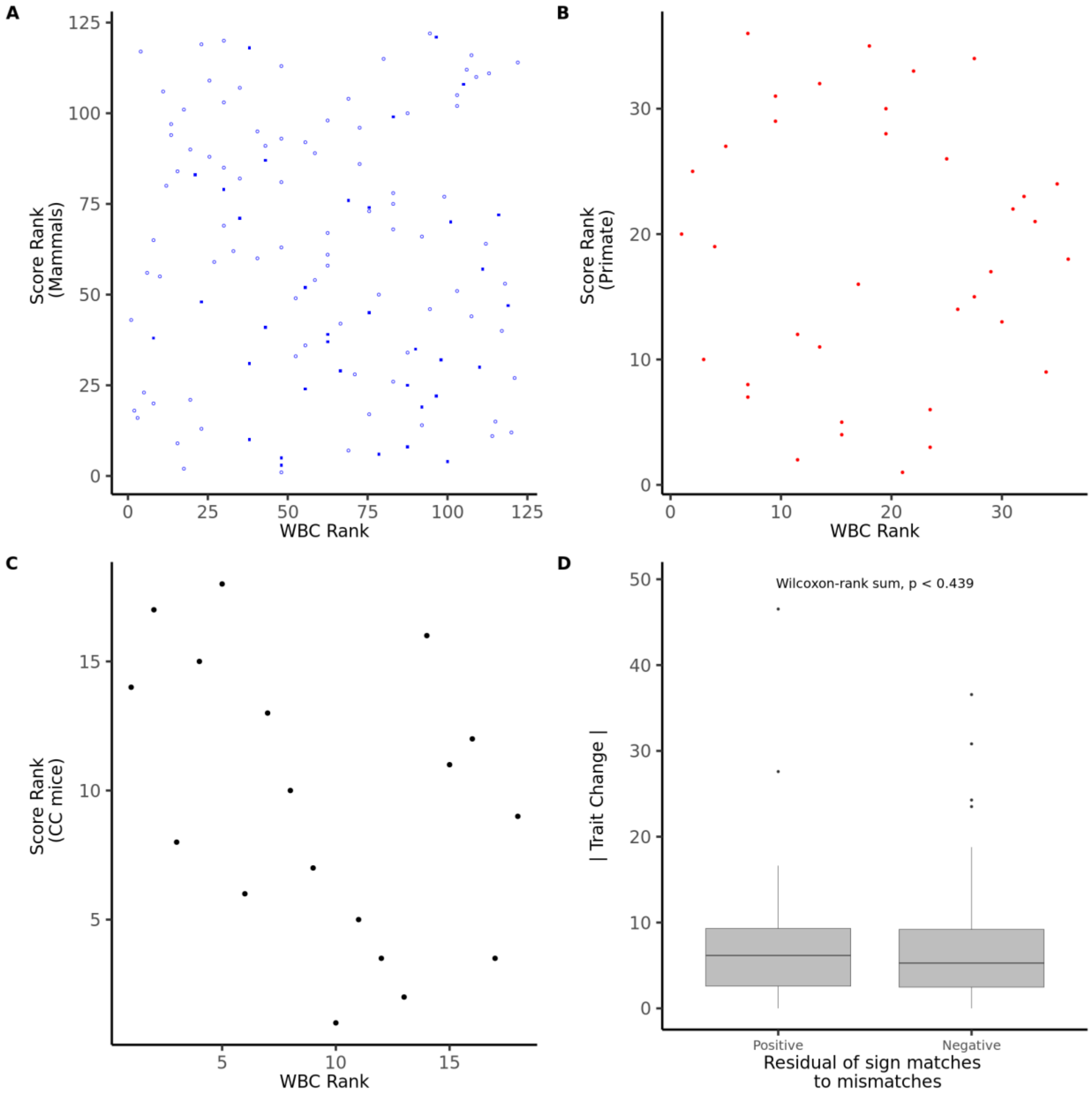


**Figure S9.** PGLS regression of white blood cell (WBC) rank against the genomic score computed using mammal conserved SNP positions (A) or primate conserved SNP positions (B). In A, closed squares represent primate species, open circle represent non-primate mammals. (C) Linear regression of white blood cell rank against genomic score computed for collaborative cross mouse lines. (D) Comparison of absolute magnitude of white blood cell change for internal branches with a positive residual of matches (allele effect direction matches phenotypic change) to mismatches compared to those with a negative change. Regression lines only show for significant relationships.

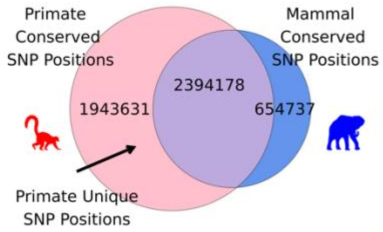


**Figure S10.** Venn diagram showing relative number of biallelic human SNP positions alignable to at least 75% of all mammals (blue circle, represented by the African elephant) or 85% of all primates (red circle, represented by the ring-tailed lemur). Positions which are uniquely alignable in primates are shown in pink. Silhouettes are from PhyloPic.

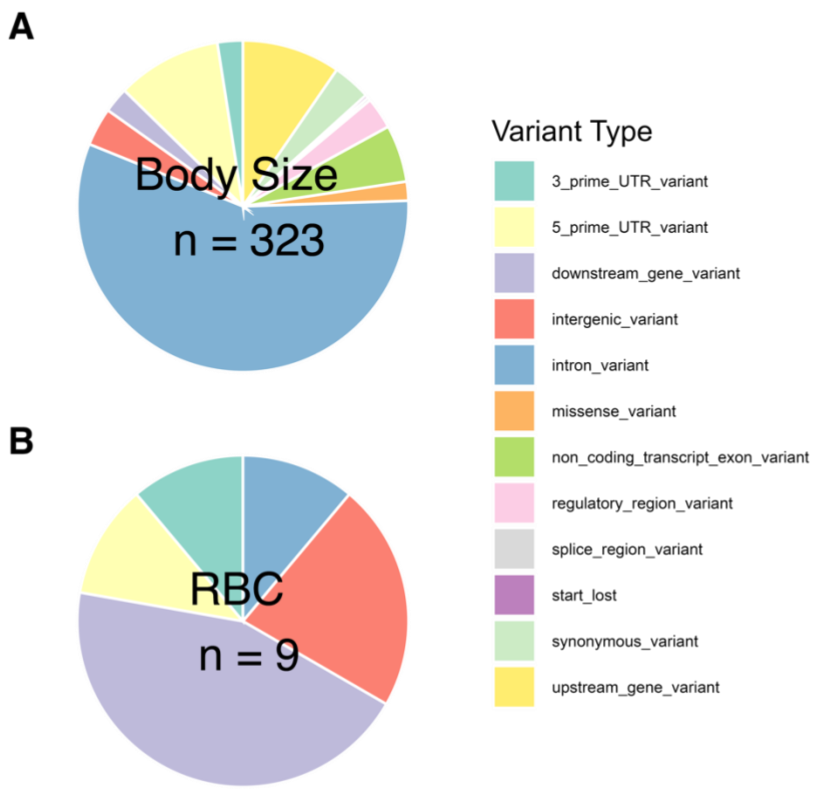


**Figure S11.** Types of putative causal variants overlapping conserved candidate *cis*-regulatory elements for body size (A) and red blood cell count (B).

### Supplementary Tables

**Table S1:** Extended test results for each trait. Includes results for the PGLS regressions for eleven traits using genomic scores calculated based on SNP positions conserved in mammals (blue), primates (red), or primates but not other mammals (pink). The number of species (n) for which genomic and phenotypic information is listed for each comparison type along with the number of GWAS significant SNPs, the sign of the relationship slope, and *P*-values determined by PGLS and after permulations. *P*-values are shown for a Wilcoxon rank-sum test of enriched trait change along internal branches with positive residuals of trait matches to mismatches and for a Spearman correlation of genomic scores and trait rank matches in eighteen collaborative cross mice strains.

**Table S2:** Candidate causative variant information for red blood count. Table include columns for (1) the human RSID (2), the trait, (3) the human genomic position (hg38), (4) the human reference and alternate alleles, (5) the direction of effect of the alternate allele, (6) the Wilcoxon P-value for the association of each allele with the trait across the mammals investigated in this study, (7) the PhyloP score for a range of 21 BPs centered around the SNP position, (8) the variant type, (9) syntenic genes within a 1 MB window centered on the SNP in human, macaque, dog, cattle, and mouse, (10) genes for which this position is a cis-eQTL based on human blood samples, and (11) predicted TF motifs binding around this SNP in both human and mouse, (13-252) aligned allele for all species used in the study which match a human allele, (253) total number of species with alignments at this position, (254) number of alignments to human reference allele, (255) number of alignments to human alternative allele, (256) *P*-value for Wilcoxon rank-sum test for phenotype rank by allelic match, (257) mean phenotype rank of species with the reference allele, (258) mean phenotype rank of species with the alternate allele, and (259) Wilcoxon test W statistic.

**Table S3:** Candidate causative variant information for body size. Table include columns for (1) the human RSID (2), the trait, (3) the human genomic position (hg38), (4) the human reference and alternate alleles, (5) the direction of effect of the alternate allele, (6) the Wilcoxon P-value for the association of each allele with the trait across the mammals investigated in this study, (7) the PhyloP score for a range of 21 BPs centered around the SNP position, (8) the variant type, (9) syntenic genes within a 1 MB window centered on the SNP in human, macaque, dog, cattle, and mouse, (10) genes for which this position is a cis-eQTL based on human blood samples, and (11) predicted TF motifs binding around this SNP in both human and mouse, (13-252) aligned allele for all species used in the study which match a human allele, (253) total number of species with alignments at this position, (254) number of alignments to human reference allele, (255) number of alignments to human alternative allele, (256) *P*-value for Wilcoxon rank-sum test for phenotype rank by allelic match, (257) mean phenotype rank of species with the reference allele, (258) mean phenotype rank of species with the alternate allele, and (259) Wilcoxon test W statistic.

**Table S4:** Resources used in this project. Lists the various datasets and computation resources used in this project and any relevant accession information.
